# Disentangling reference frames in the neural compass

**DOI:** 10.1101/2023.05.21.541641

**Authors:** Léo Dutriaux, Yangwen Xu, Nicola Sartorato, Simon Lhuillier, Roberto Bottini

**Affiliations:** Center for Mind/Brain Sciences (CIMeC); University of Trento; 38122; Trento, Italy; LaPEA; Université Gustave Eiffel; Université de Paris; F-78000; Versailles, France

**Author notes:** Corresponding authors: Léo Dutriaux, Centro Interdipartimentale Mente/Cervello (CIMeC) Università degli Studi di Trento, Via delle Regole, 101 38123 Mattarello (TN); Roberto Bottini, Centro Interdipartimentale Mente/Cervello (CIMeC) Università degli Studi di Trento, Via delle Regole, 101, 38123 Mattarello (TN).

**Keywords:** fMRI, neural compass, navigation, retrosplenial, parietal cortex, entorhinal

## Abstract

The neural system that encodes heading direction in humans is found consistently in the medial and superior parietal cortex and the entorhinal-retrosplenial circuit. However, it is still unclear whether heading direction in these different regions is represented within an allocentric or egocentric coordinate system. To investigate this problem, we first asked whether regions encoding (putatively) allocentric facing direction also encode (unambiguously) egocentric goal direction. Second, we assessed whether directional coding in these regions scaled with the preference for an allocentric perspective during everyday navigation. Before the experiment, participants learned different object maps in two geometrically similar rooms. In the MRI scanner, their task was to retrieve the egocentric position of a target object (e.g., Front, Left) relative to an imagined facing direction (e.g., North, West). Multivariate analyses showed, as predicted, that facing direction was encoded bilaterally in the superior parietal lobule (SPL), the retrosplenial complex (RSC), and the left entorhinal cortex (EC). Crucially, we found that the same voxels in the SPL and RSC also coded for egocentric goal direction. Moreover, when facing directions were expressed as egocentric bearings relative to a reference vector, activities for facing direction and egocentric direction were correlated, suggesting a common reference frame. Besides, only the left EC coded allocentric goal direction as a function of the subject’s propensity to use allocentric strategies. Altogether, these results suggest that heading direction in the superior and medial parietal cortex is mediated by an egocentric code, whereas the entorhinal cortex encodes directions according to an allocentric reference frame.

## 1. Introduction

To navigate successfully, it is crucial for an organism to know its current position and heading direction. In rodents, head direction cells are thought to constitute the neural substrates of facing direction. Indeed, these neurons discharge in relation to the organism’s facing direction with respect to the environment (Taube et al., 1990), working as a neural compass. This neural compass was further suggested to be involved in the representation of goal direction during navigation (Bicanski and Burgess, 2018; Byrne et al., 2007; Erdem and Hasselmo, 2012; Schacter et al., 2012), allowing the computation of the movements required to reach a goal from the current location and orientation.

The neural compass in humans has been mainly studied using fMRI and both univariate (adaptation) and multivariate (MVPA) approaches. These methods allowed to isolate the brain regions representing imagined heading direction in a familiar environment (e.g., the university campus) or during navigation in virtual reality (Baumann and Mattingley, 2010; Chadwick et al., 2015; Chrastil et al., 2016; Kim and Maguire, 2019; Marchette et al., 2014; Shine et al., 2019, 2016; Vass and Epstein, 2017, 2013). These studies revealed a number of brain regions representing heading direction, including in particular the entorhinal cortex, the retrosplenial complex, and superior parietal regions. Whereas some of these studies focused either on facing or goal direction (Baumann and Mattingley, 2010; Chrastil et al., 2016; Shine et al., 2019, 2016; Vass and Epstein, 2017, 2013), some others found that the same areas represented both types of heading directions (Chadwick et al., 2015; Marchette et al., 2014). Since the neural compass codes for directions relative to the environment, previous works have suggested that these regions encode directions in an allocentric reference frame, independent from the agent’s vantage point (Marchette et al., 2014; Shine et al., 2016; Vass and Epstein, 2017; Weisberg et al., 2018). However, it is still unclear whether such a directional code is encoded according to an egocentric or an allocentric reference frame. In most cases, putatively allocentric heading can indeed be accounted for by egocentric bearings to specific landmarks (Marchette et al., 2014). For instance, when entering a new environment, it is possible to choose a principal reference vector (e.g., directed to a specific landmark or from the first-perspective acquired) and to code all directions as an egocentric bearing relative to this vector (Shelton and McNamara, 2001). Hence, any putative allocentric direction can be expressed both as an allocentric heading and as the egocentric angle required to rotate from the principal reference vector to this direction (see Figure 1).

**Figure 1.**
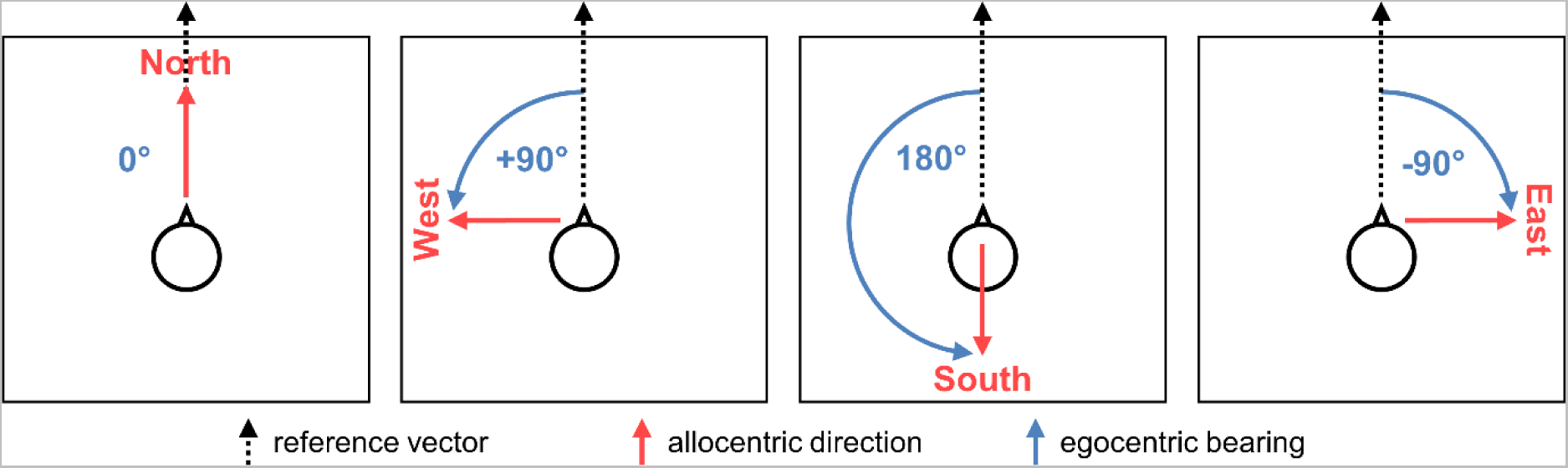
Example of how any allocentric directions (in red) can be expressed as an egocentric bearing (in blue) relative to a reference vector.

The present study aimed to disentangle these two frames of reference using fMRI and Representational Similarity Analysis (RSA) (Kriegeskorte et al., 2008). For that purpose, participants were familiarised first with two virtual rooms containing each layouts of four objects (see Figure 2A-B). They were asked to remember the location of the objects within the two virtual rooms and trained to perform the fMRI task. During scanning, each trial (see Figure 2C) started with the presentation of an orienting cue consisting of a stylized head in the middle of the room facing one of the walls. Participants were instructed to imagine themselves facing the cued wall. Afterward, a picture of an object was presented, and participants had to recall the egocentric (goal) direction of that object (left, right, back, front) given the current facing direction. We hypothesized first that heading direction is indeed encoded in the three areas of interest mentioned above: the entorhinal cortex (EC), retrosplenial complex (RSC), and superior parietal lobule (SPL). Then, since the reference frame underlying the coding of heading direction is uncertain, the present study is designed to disentangle them in two different ways. First, we investigated whether the very same regions that encoded (putatively) allocentric facing direction also encoded (non-ambiguous) egocentric goal direction. Indeed, while allocentric directions can be expressed egocentrically with respect to a principal reference vector, an egocentric direction like left, back, or right can only be formulated with respect to the current vantage point. Second, because of the intrinsic ambiguous nature of allocentric directions, we assessed whether these regions encode direction in an allocentric reference frame using an external validity method. Namely, whether a region was encoding allocentric heading direction as a function of the subject’s propensity to use allocentric navigation strategies in daily life.

**Figure 2.**
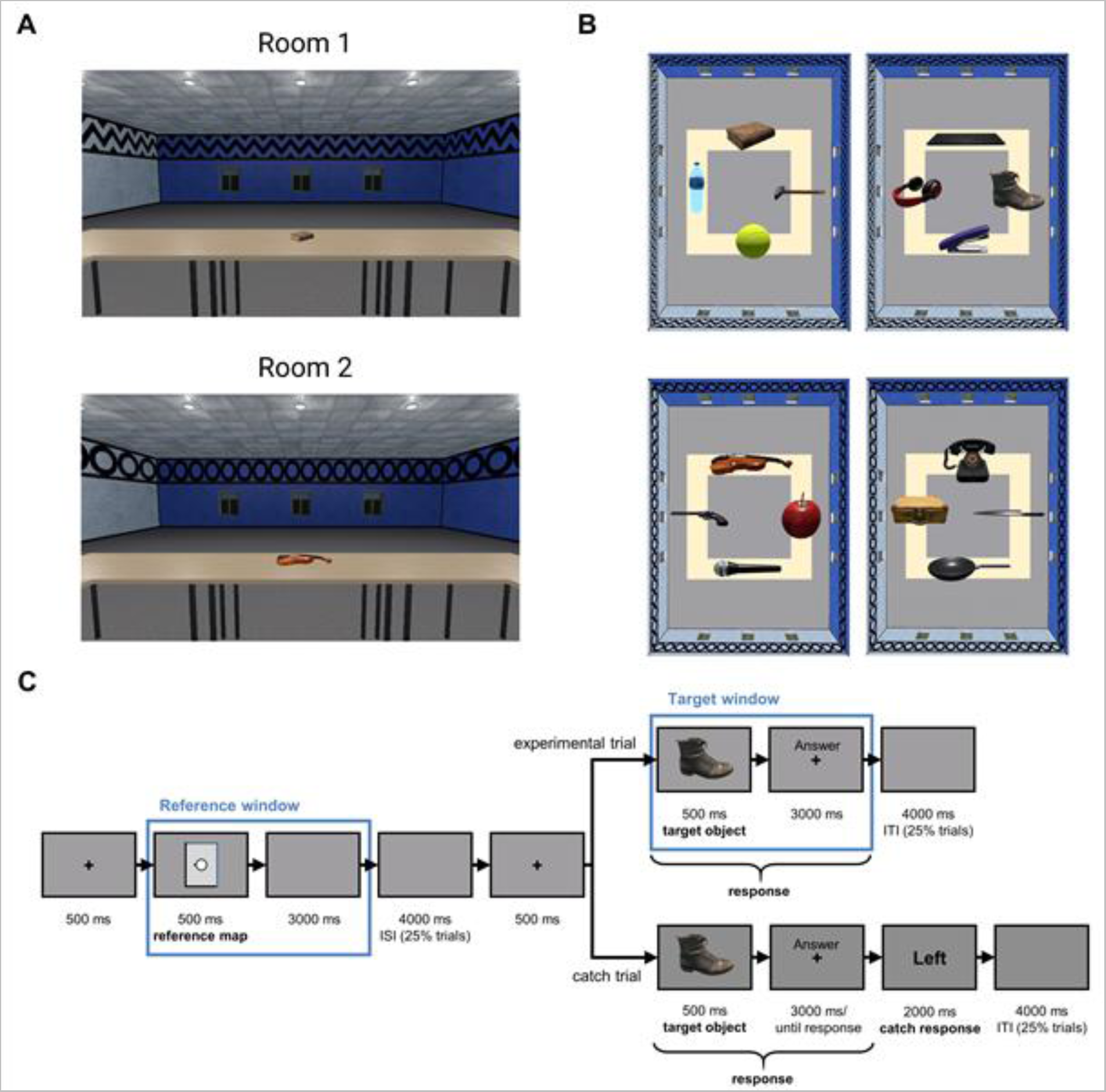
Materials. (A) Views of the two rooms. Room 1 has a zigzag pattern on the upper part of the walls, while Room 2 has circles. Participants were in the middle of the room and explored the rooms rotating their point of view clockwise or counterclockwise using the right and the left arrow, respectively. (B) Examples of two versions of each room. (C) Sequence of events of the fMRI task. ISI: Interstimulus interval; ITI intertrial interval.

## 2. Methods

### 2.1. Participants

Thirty-four right-handed native Italian speakers with no history of neurological or psychiatric disorders participated in this experiment (17 women and 17 men, mean age = 23.94, SD age = 3.90, range 18-35). The ethical committee of the University of Trento approved the experimental protocol. All participants provided informed written consent before the start of the experiment, and they received a payment as compensation for their time. We discarded one participant because of a catch trials accuracy lower than 2 SDs below the mean.

### 2.2. Materials

#### 2.2.1. 3D Rooms

Two virtual rectangular rooms (Room 1 and Room 2) were created with *Unity*. These rooms could be distinguished based on the frieze pattern on the upper part of the walls: Room 1 had a zigzag pattern, while Room 2 had circles (see Figure 2A). In these rooms, one of the two short walls and one of the two long walls were white, while the two remaining walls were blue. Hence, a wall could be recognised by its specific combination of length (short or long) and color (blue or white). The participants’ point of view was placed in the middle of the room and was stationary. Four tables were surrounding this point of view, forming a square paralleling the room’s walls, and one object was placed at the middle of each of these four tables. Two different versions of each room were further created (Room 1 version a and b; Room 2 version a and b), containing two different object layouts to dissociate object identities and object locations. In sum, in each version, the room contained a different layout with different objects. These layouts consisted of four objects assigned randomly to one of the four slots located each in front of one of the four walls (see Figure 2B).

#### 2.2.2. SDSR questionnaire

The SDSR (Sense of Direction and Spatial Representation) questionnaire allowed us to assess the participants’ propensity to use the survey perspective (i.e., bird’s-eye view representations of the environment) during spatial navigation in everyday life (Pazzaglia et al., 2000). This questionnaire comprises 11 items on a 5-point scale (except for three items that included only three alternatives). Even if the questionnaire provides several scores measuring the propensity to use several perspectives, we were primarily interested in the survey perspective, which is usually associated with an allocentric frame of reference. The survey items comprised four questions that assessed how much participants tend to use a representation “from above” (i.e., a “map-like representation”) while navigating a city. The scores could range from −2 to 14. Although the questionnaire addressed navigation in a large-scale environment, it has been validated by its authors using a pointing task in a room (Pazzaglia et al., 2000) and has been shown to be informative in other studies using smaller spaces (Lhuillier et al., 2018). Furthermore, it is known that individuals’ strategies are usually the same in large and small-scale environments (Lawton, 1996).

### 2.3. Behavioral procedures

The experiment was organised in three sessions: two training sessions – the familiarisation and the rehearsal session – and the scanning session. These sessions will now be described in detail in turn. All tasks were developed with Python 3.7 using the Neuropsydia package (Makowski and Dutriaux, 2017).

#### 2.3.1. Session 1: Familiarisation session

This first training session was conducted online, around a week before the scanning session. It was designed to familiarize participants with the rooms to allow them to construct a mental representation of the environments and to train them to perform the main task. Besides exploring the rooms, participants’ allocentric knowledge of the rooms was assessed with three tasks (see Figure S1). Task *a* aimed to assess participants’ knowledge of the spatial location of the objects relative to the walls of the room, while task *b* aimed to assess participants’ knowledge of the spatial location of the objects relative to the other objects. Finally, the test task was designed to ensure that the participant would easily perform the fMRI task.

The detailed sequence of tasks of this session is shown in Figure S2. Participants were first asked to explore both versions of one room and to perform task *a* to test their memory of this room. They had then to repeat the same procedure with the other room. Second, the participants had to perform the same sequence of exploration and memory test with task *b*. Third, after participants were given the opportunity to explore all four versions another time, they had to perform a version of tasks *a* and *b* that tested their memory of all rooms at once. The order of exploration of the rooms and the order of exploration of versions a and b of each room were counterbalanced across participants. This resulted in four possible exploration orders. Importantly, if the participant’s accuracy was lower than 85% at one of the tasks, they were asked to explore again the room’ versions related to this task and to perform it again. If needed, they had to repeat the exploration/testing sequence until they reached the threshold of 85%. The exploration phase and the three tasks will now be detailed in turn.

##### 2.3.1.1. Exploration

To familiarize the participants with the virtual environment set up, they were instructed to explore an empty room with no pattern on the wall as an example. While exploring a room, they could only perform a rotation movement of their point of view from the middle of the room by pressing the left or right arrow. No other movement inside the virtual environment was possible. The first exploration of each version of the rooms lasted two minutes, and participants were asked to memorize the objects and their position. Any later exploration lasted a maximum of one minute. When entering for the first time a version of a room, the vantage point was oriented towards the short blue wall (see Figure 2A). For simplicity, we refer to this wall as being the North wall. Consistently, we associated the other walls with their corresponding cardinal direction.

##### 2.3.1.2. Task a

As represented in Figure S1A, a trial of this task consisted of the presentation of a schematic map of a room on the left and of a target object on the right. The map was rectangular and included all information about the walls, but no information about the objects, as four white squares were placed on the map at all four possible object locations. Using the mouse, participants had to indicate with a left click the white square corresponding to the object’s location relative to the walls. Task *a* comprised 16 trials after exploring a single room, and 32 after exploring both rooms. Each object was presented as target twice in random order. The map was presented in 4 different orientations (i.e., the north oriented towards the top, the right, the bottom, or the left of the screen), and this orientation was counterbalanced across trials.

##### 2.3.1.3. Task b

Similar to task a, a trial of this task consisted of the presentation of a schematic map of a room on the left and of a target object on the right. Different from task *a* however, the walls were not displayed, and one reference object was placed at one of the four possible locations (see Figure S1B). Participants had to indicate with the mouse the target object’s location relative to the reference object. Task *b* comprised 16 trials after the exploration of a single room, and 32 after the exploration of both rooms. Each object in the room was a target twice, and objects were presented in random order. The reference object was chosen randomly, and the orientation of the map on the screen was counterbalanced.

##### 2.3.1.4. Test task

An example of a trial is presented in Figure S1C. A trial started with the presentation of a fixation cross for 2000 ms. Then, a reference map with a character facing one of the four walls was shown on the screen in one of the four possible orientations for 500 ms. At this time, participants were instructed to imagine facing the wall cued by the character on the screen. Immediately after the map, a fixation cross was displayed along with the word “Ready”. Participants had to press the space bar when they finished imagining which triggered the disappearance of the word. After 3500 ms from the outset of the map, the target object was displayed for 500 ms, followed by a 3500 ms screen prompting them to answer. They then had a total of 4000 ms to indicate the egocentric position of the object relative to their imagined heading (i.e., front, back, left, or right) using the directional arrows on the keyboard. During this task, each object was presented eight times, twice for each of the four egocentric directions. This resulted in 128 trials, which were presented randomly in 4 blocks of 32 trials. The four orientations of the reference map were counterbalanced across the eight trials of each of the 16 allocentric × egocentric levels. Therefore, each map orientation was presented twice for each of these 16 levels. In addition, each object appeared twice in a block. There was a one-minute break between each block. Only participants that scored at least 85% on this test were considered for the subsequent phases.

#### 2.3.2. Session 2: Rehearsal session

The second training session was conducted in the lab, two days before the scanning session at the earliest. After re-exploring the rooms, participants were simply asked to perform the final task of the first session. This session allowed participants to rehearse their memory of the room before the scanning session and ensured that they could still easily perform the fMRI task.

#### 2.3.3. Session 3: Scanning session

Because the fMRI task was slightly different from the test task of the training sessions, participants were first trained to perform it before entering the MRI. There were two main differences between these tasks (see Figure 2C). First, the answer in response to the reference map was no longer needed. Second, instead of indicating the egocentric location of the target objects with the directional arrows, participants were instructed to indicate when they were ready to answer. This was to avoid a confound between motoric activity and egocentric direction. Each experimental trial started with a 500 ms fixation cross. Then, a reference map was displayed for 500 ms, followed by a 3000 ms interval. At this moment, an interstimulus interval of 4000 ms was present in 25% of the trials (Zeithamova et al., 2017). Next, another fixation cross was presented for 500 ms. The target object was then displayed for 500 ms, followed by a 3000 ms interval. Participants were instructed to indicate when they were ready to answer and could do so from the beginning of the target object time window until the end of the 3000 ms interval. At this moment, an intertrial interval of 4000 ms was present in 25% of the trials. At the end of 20% of the trials, a catch trial was added to ensure that participants were actually doing the task. In these trials, an egocentric directional word (Front, Right, Back, Left) appeared on the screen for 2000 ms. Participants had to indicate whether this word matched the actual egocentric direction of the object. Fifty percent of the catch trials matched the actual direction of the object. In the experimental trials, each object was presented in all 16 conditions resulting from the combination of the four allocentric directions and the four egocentric directions. The four orientations of the reference map were counterbalanced across the 16 trials of each of the 16 allocentric × egocentric levels. Therefore, each map orientation was presented four times for each one of these 16 levels. This resulted in 256 experimental trials, to which 64 catch trials were added. These 320 trials were arranged in 8 runs of 40 trials (32 experimental trials and eight catch trials). Each block included trials for only one version per room, which means that there were two blocks for each room × version combination. Each object appeared twice in each block. Within a block, there was a catch trial every four experimental trials, placed in a random position within these four trials. This was done to spread the catch trials along the whole run. Participants were given the opportunity of a break between each run. After the fMRI session, they had to complete the SDSR scale.

### 2.4. MRI procedures

#### 2.4.1. MRI data acquisition

MRI data were acquired using a MAGNETOM Prisma 3T MR scanner (Siemens) with a 64-channel head-neck coil at the Centre for Mind/Brain Sciences, University of Trento. Functional images were acquired using the simultaneous multislice echoplanar imaging sequence (multiband factor = 5). The angle of the plane of scanning was set to 15° towards to the chest from the anterior commissure - posterior commissure plane to maximize the signal in the MTL. The phase encoding direction was from anterior to posterior, repetition time (TR) = 1000 ms, echo time (TE) = 28 ms, flip angle (FA) = 59°, field of view (FOV) = 200 mm × 200 mm, matrix size = 100 × 100, 65 axial slices, slices thickness (ST) = 2 mm, gap = 0.2 mm, voxel size = 2 × 2 × (2 + 0.2) mm. Three-dimensional T1-weighted images were acquired using the magnetization-prepared rapid gradient-echo sequence, sagittal plane, TR = 2140 ms, TE = 2.9 ms, inversion time = 950 ms, FA = 12°, FOV = 288 mm × 288 mm, matrix size = 288 × 288, 208 continuous sagittal slices, ST = 1 mm, voxel size = 1 × 1 × 1 mm. B0 fieldmap images, including the two magnitude images associated with the first and second echoes of the images and the phase-difference image, were also collected for distortion correction (TR = 768 ms, TE = 4.92 and 7.38 ms).

#### 2.4.2. fMRI Preprocessing

The preprocessing was conducted using the SPM12 for MATLAB ® (https://www.fil.ion.ucl.ac.uk/spm/software/spm12/). First, we computed each participant’s Voxel Displacement Map (VDM) using the FieldMap toolbox (Jenkinson, 2003; Jezzard and Balaban, 1995). Second, functional images in each run were realigned to the first image of the first run, and then the VDM were also coregistered to the first image and were used to resample the voxel values of the images in each run to correct for EPI distortions caused by the inhomogeneities of the static magnetic field in the vicinity of the air/tissues interface. Third, the functional images were coregistered onto the structural image in each individual’s native space with six rigid-body parameters. Lastly, a minimum spatial smoothing was applied to the functional images with a full width at half maximum (FWHM) of 2mm.

#### 2.4.3. Regions of Interests

Multiple ROI masks were used in the analysis. The entorhinal, superior parietal cortex were segmented in each subject’s native space with the Freesurfer image analysis suite 2. The entorhinal cortex masks were thresholded at a probability of 0.5, as recommended by Freesurfer 3. The location estimates for the EC were based on a cytoarchitectonic definition (Fischl et al., 2009), and masks for the SPL were based on the Destrieux atlas (Destrieux et al., 2010). Because activation in RSC in previous studies was not found in the anatomical retrosplenial cortex, we used masks of the retrosplenial cortex defined functionally as category-specific regions for scene perception (Julian et al., 2012). Anatomical RSC was defined as the combination of BA29 and 30 using MRIcron. These last masks were in MNI space and were then coregistered onto the structural image in each individual’s native space.

#### 2.4.4. First-level Analysis

Both the patterns instantiated during the appearance of the reference map and the target object were analyzed. Therefore, there were 16 experimental conditions related to the reference map (4 walls × 4 orientations) and 32 related to the target object (4 allocentric directions × 4 egocentric directions × 2 rooms). The first-level analysis was computed using the SPM12 package. The brain signal related to the reference was modeled as stick functions convolved with the canonical HRF, and the brain signal related to the target was modeled as boxcar function (duration equals to the reaction time) convolved with the canonical HRF in the time window between the presentation of the target and the response. We then used the resulting T images (48 volumes, one for each condition) in the following analyses.

#### 2.4.5. Representational Similarity Analysis

##### 2.4.5.1. RDM models

Representational Similarity Analysis (RSA) uses a correlation measure to compare a brain-based RDM obtained by calculating the pairwise correlation between patterns instantiated in all the pairs of conditions with a model-based RDM of experimental interest (Kriegeskorte et al., 2008). A brain-based RDM is a squared matrix containing the Spearman correlation distance (1−r) between two brain patterns instantiated during two different conditions. Thus, its dimension was 16 × 16 in the case of the reference window and 32 × 32 in the target window.

Accordingly, we created model-based RDMs for the reference and the target analyses. In the reference window, the facing direction model assumed that trials for which participants had to face the same wall were similar, regardless of the room’s orientation relative to the screen (see Figure 3A). In the target window, we created two model RDMs. In the facing direction model, only conditions with the same facing direction *and* the same room were considered similar. In the second RDM, the facing-generalized model, facing directions were considered similar regardless of the room (see Figure 4A).

**Figure 3.**
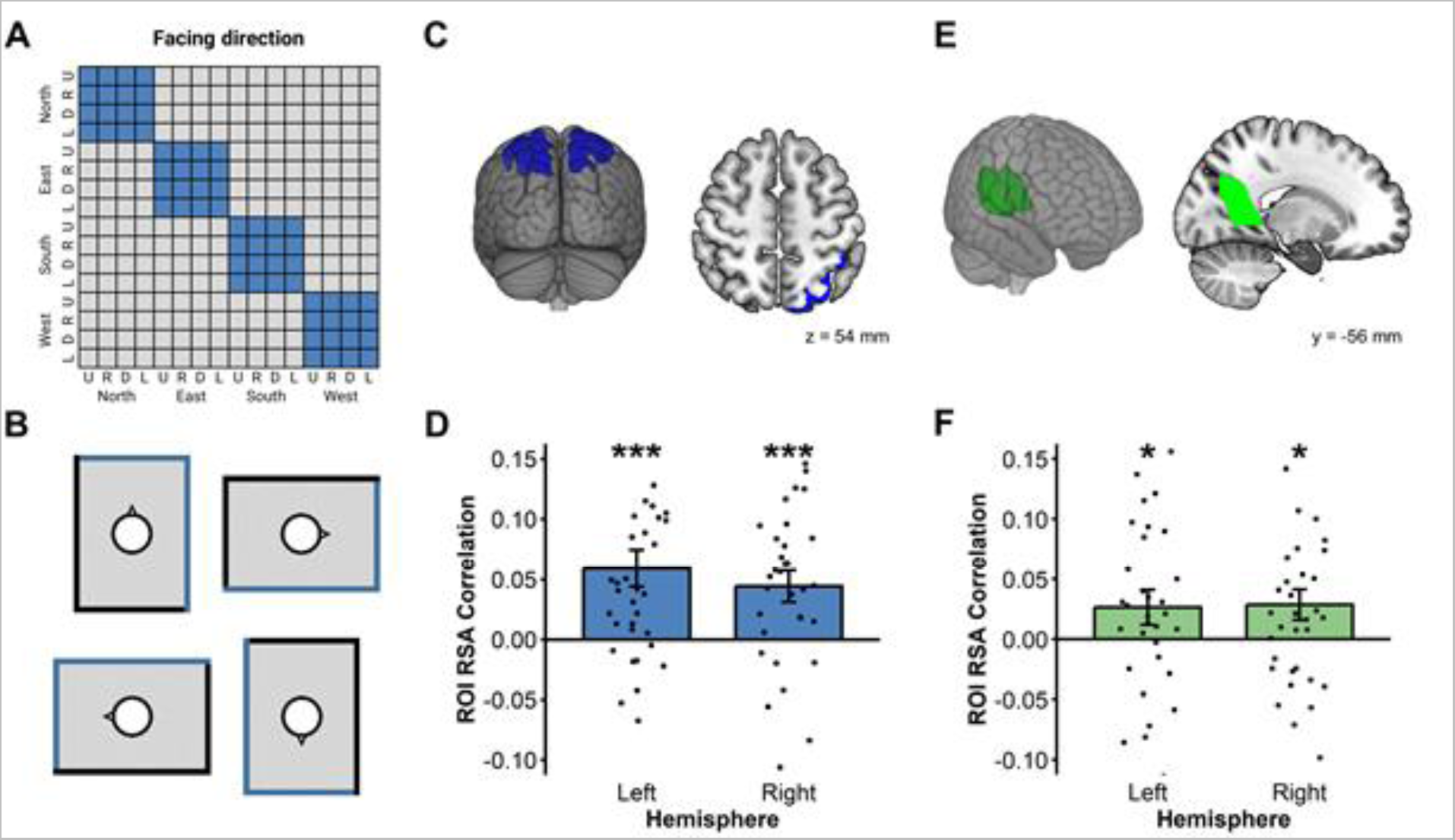
(A) The 16 x 16 RDM for reference direction, where two trials were considered similar when they shared the same facing direction, regardless of the orientation of the North on the screen (U = Up, R = Right, D = Down, L = Left). (B) For instance, in the example in the right panel, all these reference maps were cuing the same facing direction. (C) SPL ROIs (D) Both SPL showed reliable facing direction coding in the reference window. (E) RSC ROIs (F) Both RSC showed reliable facing direction coding in the reference window. For each RDM, a diamond represents the mean correlation; a box and whisker plot represent the median and inter-quartile range. (* p < .05; *** p < .001).

**Figure 4.**
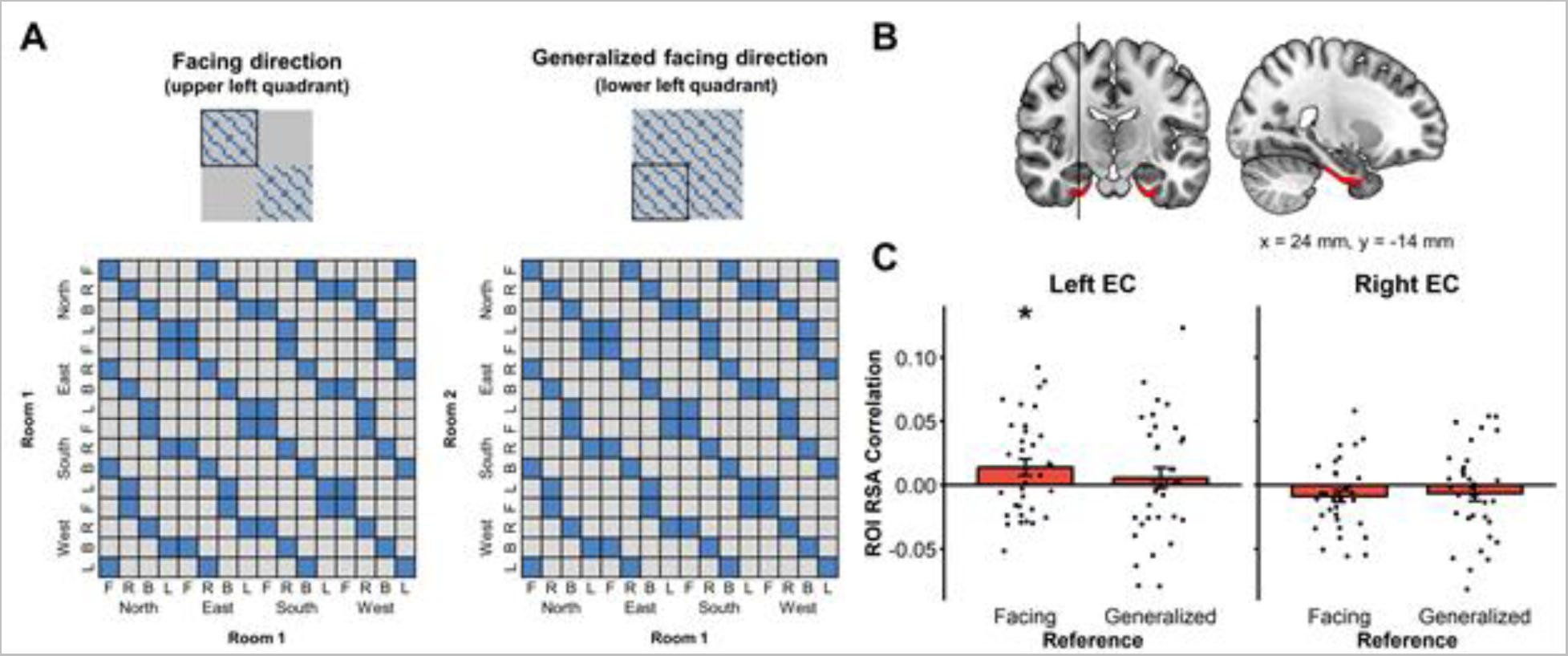
(A) In the upper panels are presented the whole 32 × 32 RDMs, including all Rooms × Allocentric directions × Egocentric directions conditions. The black square indicates which quadrant of the matrix is represented in the lower panel. The key difference between the facing and the facing-generalized model is that in the facing-generalized model, two facing directions were considered similar even if they were in different rooms (lower-right panel); while in the facing model, they were considered similar only if they were in the same room (lower-left panel) (F = Front, R = Right, B = Back, L = Left). (C) EC ROIs. (B) Left EC showed facing direction coding in the target window. For each RDM, a diamond represents the mean correlation; a box and whisker plot represent the median and inter-quartile range. (* p < .05).

We created three RDMs to disentangle between egocentric and allocentric goal direction in the target window (see Figure 5). In the egocentric model, conditions in which the target object was in the same egocentric position (e.g., to the left) were considered similar. In the allocentric model, only conditions in which the target object was placed in the same allocentric direction *and* in the same room were considered similar. Lastly, in the allocentric-generalized model, conditions in which the target object was placed in the same allocentric position independently of the room were considered similar. This last RDM was designed to test whether allocentric goal direction coding generalized across rooms with identical geometrical layouts. To control for response time variability, we also created a RDM where similarity between each condition was computed as the difference between their respective mean response times.

**Figure 5.**
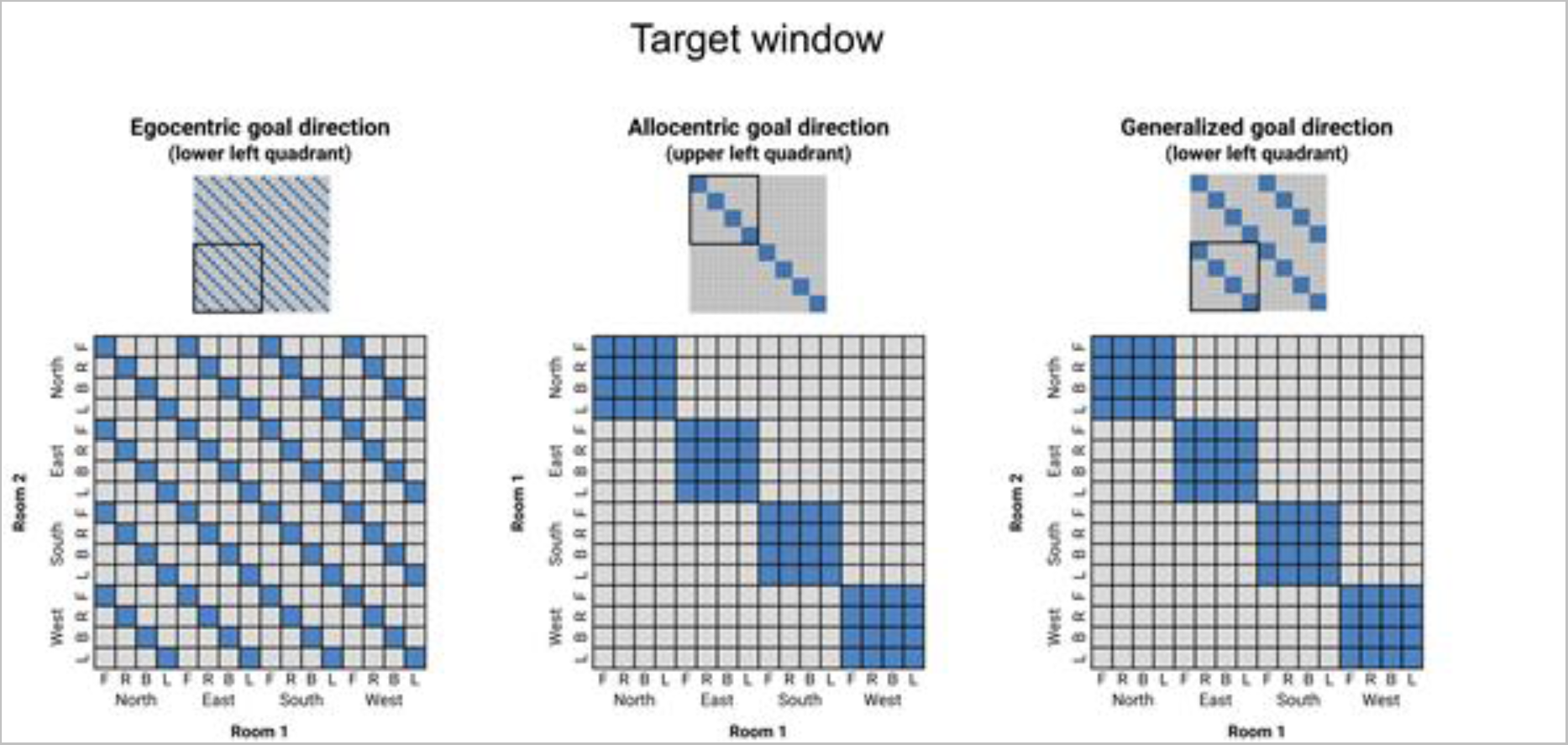
Model RDMs for target direction. In the upper panels are presented the whole 32 × 32 RDMs, including all Rooms × Allocentric directions × Egocentric directions conditions. The black square indicates which quadrant of the matrix is represented in the lower panel. The key difference between the allocentric and the allocentric-generalized model is that in the allocentric-generalized model, two objects sharing the same allocentric direction are considered similar even if they are in different rooms (right panel); while in the allocentric model, they are considered similar only if they are in the same room (middle panel). (F = Front, R = Right, B = Back, L = Left).

##### 2.4.5.2. ROI-based RSA

In the ROI-based RSA, the brain-based RDM was computed using the activity pattern of all voxels inside a given ROI. The second order-correlations between the brain-based RDM of this ROI and each model-based RDM were then performed with a partial Pearson correlation method. Partial correlations were used to regress out confounds. For example, when an effect was significant when computing the egocentric correlation, the allocentric matrix was then regressed out to obtain the correlation with the egocentric matrix controlling for the allocentric matrix. Conversely, the egocentric matrix was regressed out when the facing, allocentric, and allocentric generalized correlations were computed. This was done to measure the desired effects selectively. The resulting correlations were then tested to be greater than zero with a one-tailed one-sample t-test. To control for potential confounding effects of the RT, we used the duration modulation method by convolving each trial with a boxcar equal to the length of the trial’s RT for each participant (Grinband et al., 2008). Moreover partial correlation using the RTs RDM were implemented as a further control in some analyses.

To investigate whether individual RSA results were modulated by the participants’ propensity to use an egocentric or allocentric perspective, we computed the correlation between the individual second-order correlations for a given ROI and the individual SDSR scores. The resulting correlations were then tested to be greater than zero with a one-tailed one-sample t-test.

##### 2.4.5.3. Searchlight-based RSA

In the Searchlight RSA analysis, a brain-based RDM was calculated for each voxel using the pattern instantiated in the neighbourhood of the voxel of interest within a 6 mm sphere. After calculating the brain-based RDM, we computed the second-order correlations with each RDM model using a partial Pearson correlation method. Similar to the ROI-based RSA, egocentric RDM was regressed out for second-order correlations with allocentric RDMs, and vice versa. These second-order correlations were fisher z transformed to be used in the second-level analysis.

After computing the Searchlight images for each participant, they were normalized using the unified segmentation method and then smoothed with a gaussian kernel (FWHM of 6 mm) using SPM12. These normalized images were the input of the second-level analysis, which was performed with SnPM 13 (http://warwick.ac.uk/snpm) using the permutation-based nonparametric method (Nichols and Holmes, 2003). No variance smoothing was applied, and 10,000 permutations were performed. A conventional cluster-extent-based inference threshold was used (voxel level at p < 0.001; cluster-extent FWE p < 0.05).

To investigate whether individual differences in allocentric strategy modulated the whole-brain activity, in the second-level analysis general linear models, we used the survey score to predicit the correlation between the brain-based RDM and each model-based RDM. The resulting to a T-score volume for each model-based RDM allowed us to assess where the correlation with the model-based RDM was modulated by an individual’s propensity to use an allocentric perspective.

## 3. Results

### 3.1. Facing direction coding in the reference window is present in SPL and RSC

In the reference window, the facing direction model assumed that trials for which participants had to face the same wall were similar, regardless of the room’s orientation relative to the screen (see Figure 3A-B). ROI analyses revealed first a significant facing direction coding during the reference window in bilateral SPL and RSC (see Figure 3C-F; lSPL: t(33) = 3.90, p < .001; rSPL: t(33) = 3.34, p = .001; lRSC: t(33) = 1.80, p < .05; rRSC: t(33) = 2.26, p < .05), but not in EC (All ps > .05). Whole-brain analysis (whole-brain inferential statistics are computed with primary voxel-level p < .001, cluster-level FWE corrected p < .05) revealed an additional bilateral activation in the occipital place area (OPA; MNI coordinate of the left peak: [−38, −80, 28], t(33) = 4.46, p_FWE_ < .05; MNI coordinate of the right peak: [38, −76, 28], t(33) = 5.82, p_FWE_ = .004; see Figure S3). This effect is consistent with previous findings showing that OPA represents environmental boundaries (Julian et al., 2016).

### 3.2. Facing direction coding is present in the left EC during the target window

We then investigated the encoding of facing direction in the target window. To solve the task, participants had to keep in memory the current facing direction cued in the reference window until the target object appeared (i.e., the target window). Because the target object was presented at this moment, only then could participants encode facing direction in a room- or map-specific way. Thus, for this analysis, we created two model RDMs. In the facing direction model, only conditions with the same facing direction *and* the same room were considered similar. In the second RDM, the facing-generalized model, facing directions were considered similar regardless of the room (see Figure 4A). A significant correlation with room-specific facing direction was observed in the left EC (t(33) = 2.12, p < .05; throughout the paper, p-values are corrected for multiple comparisons across the two hemispheres; Figure 4B-C). No other ROI demonstrated room-specific or generalized facing direction coding during the target window (All ps > .05). Whole-brain analysis did not yield any significant clusters in this case.

### 3.3. The parietal and retrosplenial cortex code for egocentric but not allocentric goal direction

We created three RDMs to disentangle between egocentric and allocentric goal direction in the target window (see Figure 5). In the egocentric model, conditions in which the target object was in the same egocentric position (e.g., to the left) were considered similar. In the allocentric model, only conditions in which the target object was placed in the same allocentric direction *and* in the same room were considered similar. Lastly, in the allocentric-generalized model, conditions in which the target object was placed in the same allocentric position independently of the room were considered similar. This last RDM was designed to test whether allocentric goal direction coding generalized across rooms with identical geometrical layouts.

ROI analyses revealed a strong egocentric bilateral coding in both SPL and RSC (see Figure S5A-B; lSPL: t(33) = 7.52, p < .001; rSPL: t(33) = 6.27, p < .001; lRSC: t(33) = 5.25, p < .001; rRSC: t(33) = 3.23, p = .001), but no allocentric coding (All ps > .05). No correlations were found with the SDSR scores in these ROIs, suggesting that spatial coding in the parietal cortex did not change as a function of the propensity for a particular reference frame. Notably, this effect was also significant when we excluded the “front” condition (which, contrary to other conditions, did not require reorientation) and control for RTs (see Figure S5).

Whole-brain searchlight RSA confirmed that the parietal cortex overall coded for egocentric goal direction (Figure S4C), showing a very large bilateral clusters with a peak in the left AG (peak voxel MNI coordinates: [−48, −62, 44], t(33) = 8.17, p_FWE_ < .001) extending in the left hemisphere to the superior parietal lobule, the precuneus, and also ventrally in the inferior part of the occipitotemporal cortex (BA 37). It also spread in the right hemisphere to the AG, superior parietal lobule, and precuneus. Further, two clusters were found bilaterally in the dorsal premotor area (BA 6; left peak: t(33) = 7.56, p_FWE_ = .002; right peak: t(33) = 8.98, p_FWE_ = .004). Other clusters included the right and left posterior middle frontal gyrus (left: t(33) = 4.96, p_FWE_ < .01; right: t(33) = 5.54, p_FWE_ < .01), the left posterior cingulate cortex (t(33) = 6.80, p_FWE_ < .01), and the left pars triangularis (t(33) = 4.51, p_FWE_ < .01). (see Table S1 for details).

Next, we wanted to check whether the same voxels coding for facing direction in the reference window also coded for egocentric goal direction in the target window. For that purpose, we used the whole-brain activation maps at a lower threshold to extract four masks corresponding to the bilateral SPL and RSC clusters sensitive to facing direction during the reference window (see Figure 6A and 6B). We then used these masks to conduct ROI analyses of the egocentric and allocentric goal direction. We found that the voxels coding for facing direction during the reference window in the SPL and the RSC also coded for egocentric goal direction in the target window (lSPL: t(33) = 3.46, p < .001; rSPL: t(33) = 4.87, p < .001; lRSC: t(33) = 3.08, p = .002; rRSC: t(33) = 2.88, p = .004).

**Figure 6.**
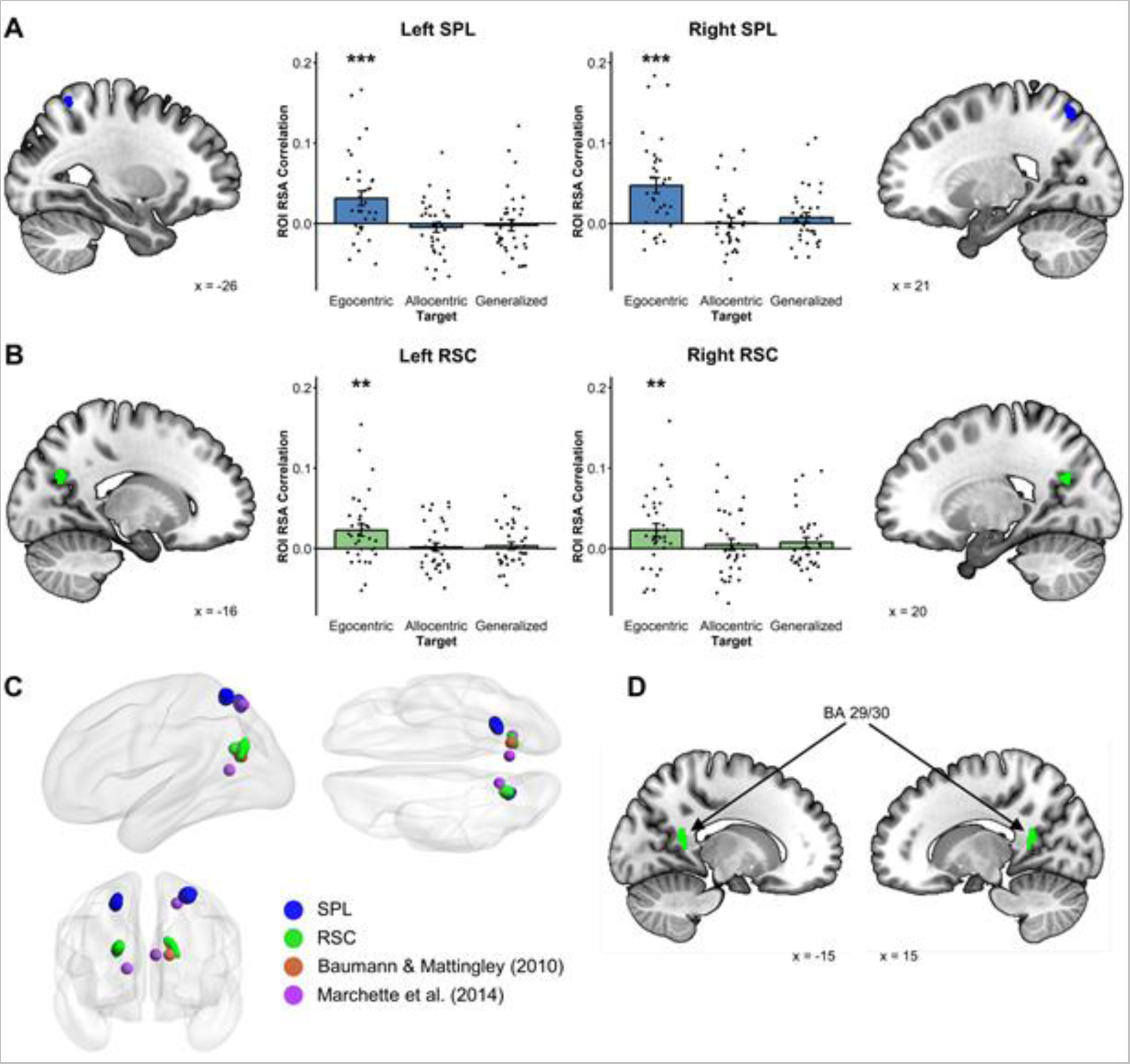
RSA Results for egocentric goal direction. (A) Voxels showing the coding of facing direction in the reference window in both SPL (extracted at p < .001) showed reliable egocentric goal direction coding in the target window. (B) Voxels showing the coding of facing direction in the reference window in both RSC (extracted at p < .05 for left RSC and p < .005 for right RSC) showed reliable egocentric goal direction coding in the target window. (C) Comparisons of the location of the RSC and SPL clusters extracted from the reference window with the peak activation coordinates in Baumann & Mattingley (2010) and Marchette et al. (2014). (D) BA29/30 Masks used in the complementary analyses. (** p < .01, *** p < .001).

Then, we compared the exact coordinate of our brain activations with those reported in other studies observing putatively allocentric facing direction in the superior parietal lobule (Marchette et al., 2014) and the retrosplenial complex (Baumann and Mattingley, 2010; Marchette et al., 2014). Our activation in the SPL overlaps with the one previously reported by Marchette and colleagues (Marchette et al., 2014), and one of the masks in the retrosplenial complex overlaps with the peak of activity reported by Baumann and Mattingley (Baumann and Mattingley, 2010) (Figure 6C). These results substantiate the comparability of our results with previous studies reporting putatively allocentric heading direction signals. However, our RSC masks were more lateral than the RSC activity reported by Marchette and colleagues. Indeed, the functionally defined RSC used here as ROI mask (see (Julian et al., 2012) and method section) comprises a large portion of the medial parietal lobe, and different studies have reported different exact functional localization of the retrosplenial cortex (Baumann and Mattingley, 2010; Marchette et al., 2014; Vass and Epstein, 2017). In some studies (Baumann and Mattingley, 2010), however, heading direction coding has been reported in the anatomically defined RSC (BA 29/30), which is outside the functional RSC mask used in ours and many other studies (Baumann and Mattingley, 2010). In an exploratory analysis, we tested whether our results generalize to this region of interest. We found facing direction in the reference window was encoded in BA 29/30 (see Figure 6D for representations of the ROIs), in the left hemisphere (t(33) = 2.25, p < .05; Corrected for multiple comparisons across hemispheres). Crucially, the same region also encoded egocentric goal direction in the target window (t(33) = 1.81, p = .04).

In sum, we performed a series of analyses using three different types of ROIs: predefined masks of the RSC and the SPL, functionally defined masks encoding facing direction in the reference window, and anatomical masks of the RSC proper (BA 29/30). In all these cases, regions encoding putatively allocentric facing direction in the reference window also encode unambiguously egocentric goal direction in the target window.

### 3.4. SPL and RSC encode both facing and goal directions relative to a principal reference vector

The results presented above suggested that the SPL and the RSC code heading direction in an egocentric fashion. One possibility is that these areas computed both facing and goal direction through the egocentric bearing relative to a principal reference vector. In the case of goal direction, this reference vector would be naturally the current imagined facing direction. Concerning the facing direction (reference window), it is known that the first experienced vantage point in a new environment tends to be used as a reference vector from which bearings are computed (Shelton and McNamara, 2001). In the present experiment, this vantage point is in the direction of what we call North in the article, which is the short blue wall (which has never been referred to as “North” to the participants). It is then possible that, in the SPL and RSC, both facing (reference window) and egocentric goal directions (target window) are computed egocentrically from a given reference vector. If that is the case, the representation of the facing direction “North” should be similar to that of the egocentric goal direction “Front”. Consequently, we should expect the following similarity pattern between the reference and the target window: North = Front, South = Back, East = Right, and West = Left.

To explore this idea, we ran a new ROI-based RSA in which we computed, for each participant, the pattern similarity between the activity for reference directions and egocentric goal directions in our ROIs. This resulted in a 4 x 4 matrix (see Figure 7A) where the North, East, South, and West reference directions on one side matched the Front, Right, Back, and Left egocentric target directions on the other side. Following the hypothesis of the principal reference vector, we expected higher average pattern similarity between matching directions (on the diagonal: North-Front, East-Right, South-Back, and West-Left) than between non-matching directions (off-diagonal). Because we wanted to see whether voxels coding for reference direction were coding similarly egocentric target direction, we used the brain masks that we extracted in the previous analyses of the reference window (Figure 6A-B). It is important to note that results are very similar when the a priori anatomical/functional ROIs are used instead. Consistent with our hypothesis, average pattern similarity is higher when directions are matching than when they are not in all parietal areas (see Figure 7B; lSPL: t(33) = 1.86, p = .03; rSPL: t(33) = 4.69, p < .001; lRSC: t(33) = 2.43, p = .01; rRSC: t(33) = 2.13, p = .02). Importantly, we did not observe this effect in the left EC (t(33) = 0.32, p = .38). These results suggest that the same egocentric representation, anchored to a specific vantage point (North in the reference window and Front in the target window) is at the basis of facing-direction and egocentric goal direction encoding in the SPL and RSC.

**Figure 7.**
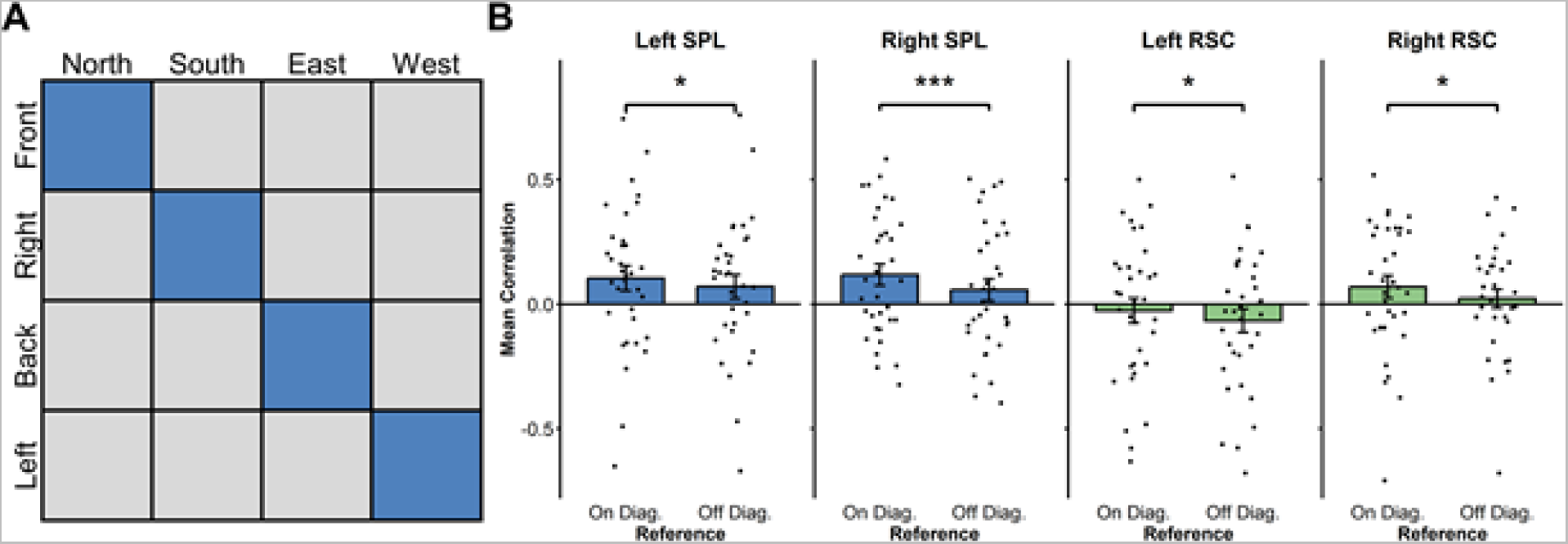
(A) Brain RDMs used to test the hypothesis that North in the reference window and Front in the target window are both used as a principal reference vector. This means that, across reference and target windows, Front is considered as matching North, while Right matches East, Back matches South, and Left matches West. In this analysis, we averaged the correlations of matching directions (in red) and the unmatching conditions (in blue) for each participant to compare these average correlations across participants. (B) RSA results for the comparison between on diagonal and of diagonal reference and egocentric target directions in SPL and RSC. Results all showed a more positive average correlation for matching conditions (* p < .05, *** p < .001).

### 3.5. Participants’ propensity for allocentric perspective modulates goal-direction coding in the EC

The ROI analysis did not yield any reliable group-level allocentric or allocentric-generalized goal direction coding either in the EC or the parietal ROIs (see Figures S6B). On the other hand, we observed a significant modulation of the allocentric and allocentric-generalized coding in the left EC by the allocentric (survey) score measured with the SDSR questionnaire (see Figure 8B-D; allocentric: r = .33, t(32) = 1.98, p < .05; allocentric-generalized: r = .38, t(32) = 2.32, p = .01). This suggests that, in our experiment, the allocentric coding in the left EC depends on participants’ propensity to use an allocentric perspective during everyday navigation.

**Figure 8.**
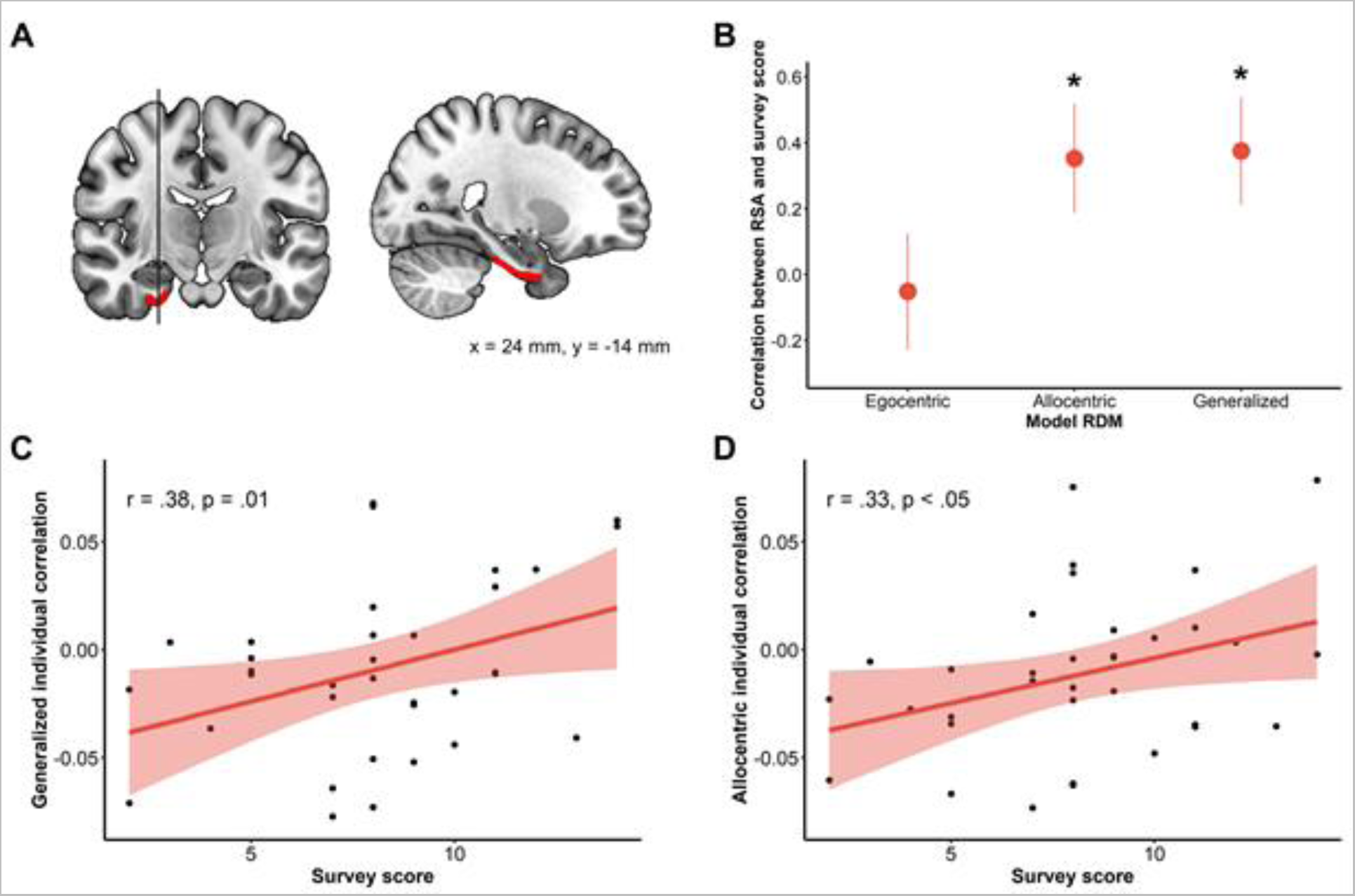
Allocentric coding in the left EC is modulated by the participant’s propensity for the allocentric perspective. (A) Left EC ROI. (B) Correlations between the ROI results in the left EC and the survey score (* p < .05). (C) Scatterplot of the correlation between the allocentric coding in the left EC and the individual score at the survey scale of the SDSR. (D) Scatterplot of the correlation between the allocentric-generalized coding in the left EC and the individual scores at the survey scale of the SDSR.

The whole-brain analysis led exclusively to bilateral occipital V1 activations (Figure S6C), both with the allocentric (left: t(33) = 8.36, p_FWE_ < .001; right: t(33) = 5.54, p_FWE_ < .001) and the allocentric-generalized model (left: [−20, −98, 12], t(33) = 5.23, p_FWE_ = .003; right: t(33) = 5.27, p_FWE_ = .002) (see Table S2 for details). This was likely due either to the reactivation of the visual information related to the wall or to the fact that, in each allocentric direction, the same objects are presented several times throughout the trials (although different objects appeared in the same allocentric direction). Besides, contrary to the left EC, activity in V1 was not correlated to the propensity to use an allocentric reference in everyday life (All ps > .10).

## 4. Discussion

The reference frame underlying the representation of heading direction in different parts of the brain remains largely ambiguous. Although previous studies found that the entorhinal cortex (EC), the retrosplenial cortex (RSC), and the superior parietal lobule (SPL) coded for facing and/or goal direction, they generally did not enable the disentanglement of egocentric and allocentric reference frames. The present study used a reorientation task which allowed us to address this question by testing (i) whether the same regions that encoded (putatively) allocentric facing direction also encoded (unambiguously) egocentric goal direction, and (ii) whether the activity in these regions was modulated by the subject’s propensity to use allocentric strategies in daily life. Results confirmed first that the EC, the RSC, and the SPL all represent facing direction. Up to that point, this effect could result from both allocentric and egocentric processing. However, we found that RSC and the SPL also encoded egocentric goal direction (whether an object is on the left/right/front/back independently of its position in the map), a result that could not emerge from an allocentric coding. This result raises the possibility that these regions actually represent heading direction according to an egocentric reference frame. On the other hand, the EC did not demonstrate any egocentric coding, and allocentric goal direction coding in this region was uniquely modulated by participants’ propensity for an allocentric perspective. Thus, in agreement with previous findings (Chadwick et al., 2015; Shine et al., 2019), the entorhinal cortex seems to encode heading direction in an allocentric reference frame. Overall, these results suggest that the neural compass can operate within different reference frames in different brain regions.

The present study replicated the results of previous studies finding the involvement of the EC, RSC, and SPL in facing direction coding (Baumann and Mattingley, 2010; Chadwick et al., 2015; Marchette et al., 2014; Vass and Epstein, 2017, 2013). Our finding that the EC heading direction system seems to operate within an allocentric reference frame is in keeping with the hypothesis that the fMRI signal is driven, at least in part, by the activity of head-direction cells (Taube et al., 1990). However, the fact that an egocentric reference frame provides a better account for the heading-related activity in medial and superior parietal cortices suggests that the neural compass in these regions arises from a different neural mechanism than the allocentric direction coded by HD cells. One possibility is that the neural activity observed in SPL and RSC comes from hypothetical reference vector cells, which would code for the egocentric bearing relative to a principal reference vector (Marchette et al., 2014). For instance, during the reference window, participants may take one of the walls as the principal reference vector (Shelton and McNamara, 2001) and compute the facing direction egocentrically in reference to that wall. In the target window, the current facing direction (the Front direction) could be defined as the new principal vector, and all directions would then be coded as an egocentric bearing from this principal reference vector. The analysis of the similarity between the brain activity across the reference and the target window backed this idea. Indeed, in both SPL and RSC, we observed higher average correlations between directions that matched according to the reference-vector model (North = Front, East = Right, South = Back, and West = Left) than between non-matching directions. These findings suggest that, in the SPL and RSC, an egocentric representation anchored to a specific direction (North or Front) is used to guide re-orientation for both facing and egocentric goal direction. In line with these results, a previous study that used a re-orientation task in a larger natural environment (a university campus) showed that putatively allocentric heading directions (North, South, East, West) were encoded in RSC both when the starting point and the target buildings were indicated with realistic pictures and when they were conveyed verbally. However, when the similarity between brain activity in RSC was compared across the two tasks (visual and verbal), only the North heading direction showed a similar pattern across conditions (Vass and Epstein, 2017). Vass and colleagues hypothesized that the RSC preference to represent north-facing headings arose because the RSC represents environments according to a particular reference direction (McNamara et al., 2003). Besides, they could not establish whether heading directions relative to that reference direction were computed egocentrically or allocentrically. In our experiment, not only we provide evidence that, in the RSC and SPL, heading is derived relative to a reference vector, but also that this computation is done within an egocentric frame of reference.

The present results showed that the representation of allocentric goal direction in the left EC was modulated by participants’ propensity for the allocentric perspective in everyday life. This, together with the presence of facing direction coding and the absence of egocentric coding, suggests that the EC coded for heading direction in an allocentric frame of reference (see also Chadwick et al., 2015). Consistently, the entorhinal cortex has strongly been associated with allocentric representation in the literature, particularly through the presence of grid cells (Hafting et al., 2005), which are thought to provide the scaffolding of allocentric representations (Buzsáki and Moser, 2013). Contrary to previous results (Chadwick et al., 2015; Shine et al., 2019), we did not find a consistent representation of allocentric goal direction in the entorhinal cortex across subjects (i.e., independently from their everyday navigation style). One possible reason for this discrepancy is that we did not explicitly ask subjects to provide the allocentric location of the target object (North, South, East, West) during the task, but only the egocentric one (Front, Back, Right, Left). Thus, participants could solve the task relying solely on egocentric information. Our result suggests that the activation of an allocentric map to retrieve the position of objects is not automatic. This interpretation is in line with previous studies showing that different cognitive styles in spatial strategies lead to the activation of partially different neural networks during the same spatial task (Iaria et al., 2003; Jordan et al., 2004). We might have failed to observe allocentric goal direction coding in the RSC for similar reasons. Indeed, according to a prominent spatial memory model (Bicanski and Burgess, 2018; Byrne et al., 2007), the RSC should serve as a hub where spatial information is transformed across reference frames. If that is the case, one should expect to find both allocentric and egocentric goal direction coding in this region. Nevertheless, if the activation of an allocentric map is indeed not necessary for the task, reference frames transformation might not have been necessary either.

## 5. Conclusion

Overall, the present work allowed to disentangle between different reference frames supporting the representation of heading direction across different brain regions. We showed that superior and medial parietal regions encode not only facing direction, as already suggested in previous studies (Baumann and Mattingley, 2010; Marchette et al., 2014; Vass and Epstein, 2017), but also egocentric goal direction. This finding suggests the use of a common egocentric reference frame to represent heading direction in these areas. On the other hand, no egocentric coding emerged in the entorhinal cortex, which, beyond representing facing direction, also represents allocentric goal direction as a function of the individual propensity to use allocentric navigational strategies in everyday life. Although limited to a particular spatial setting (small environments without translation or actual head rotation of the observer; Shine et al., 2016), our study highlights the necessity to investigate how different brain regions may encode similar spatial features by mean of different computations across reference frames. Beyond space, one can wonder whether the same sort of mechanism would apply in non-spatial domains. Indeed, there is now evidence that the EC and the PC can also generalise across non-spatial domains, particularly conceptual domains (Bottini and Doeller, 2020). Future work should focus on whether the same low-dimensional geometries relying on the same brain areas underlie the structuring of conceptual domains across different reference frames.

## Data and code availability statement

Our code is publicly available at https://github.com/BottiniLab/allo-ego, and data is available from the corresponding author upon request, without restriction.

## Acknowledgement

We thank Simone Viganò for suggestions on the early draft.

## Author contributions

**Léo Dutriaux:** Conceptualization, Methodology, Software, Formal analysis, Investigation, Writing - Original Draft, Visualization. **Yangwen Xu:** Conceptualization, Methodology, Formal analysis, Writing - Original Draft. **Nicola Sartorato:** Formal analysis, Investigation, Writing - Original Draft, Visualization. **Simon Lhuillier:** Methodology, Software, Writing - Review & Editing, Visualization. **Roberto Bottini:** Conceptualization, Methodology, Resources, Writing - Original Draft, Visualization, Supervision, Funding acquisition.

