## Supplemental materials for "Disentangling reference frames in the neural compass"

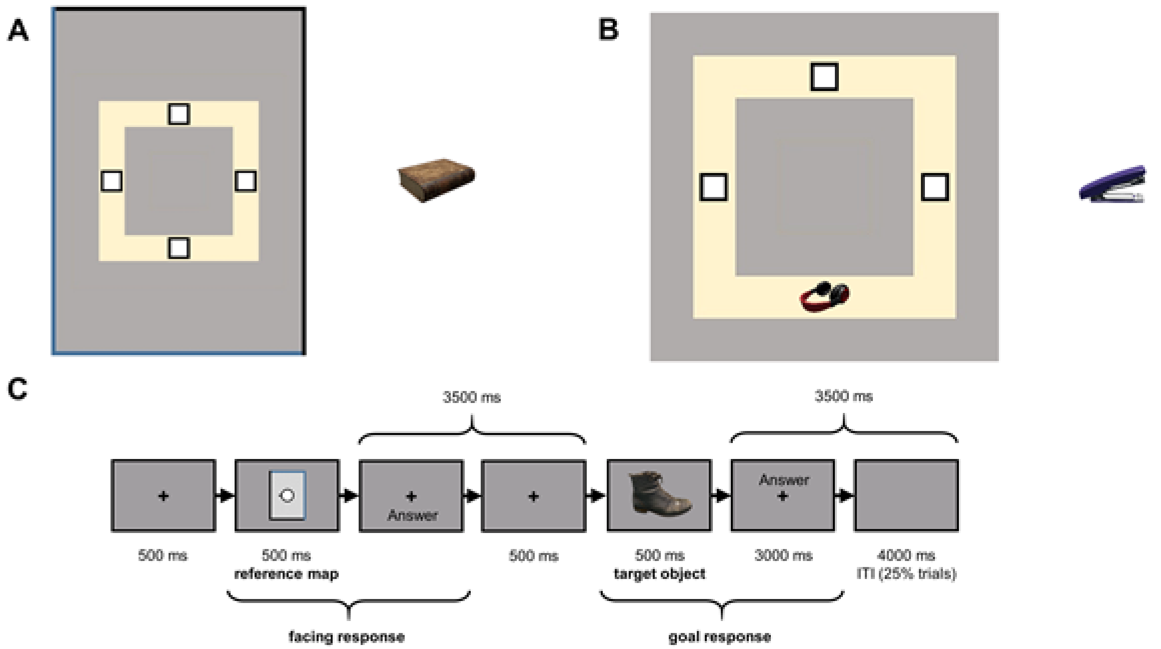


***Figure S1.*** *Tasks of the training sessions. (A) Task a required participant to click on the right location considering the orientation of the room indicated by the walls. (B) Task b required participants to click on the right location considering the position of a reference object already placed on the table. (C) The sequence of events of the test phase in sessions 2 and 3 that were designed to train participants to perform the fMRI task.*

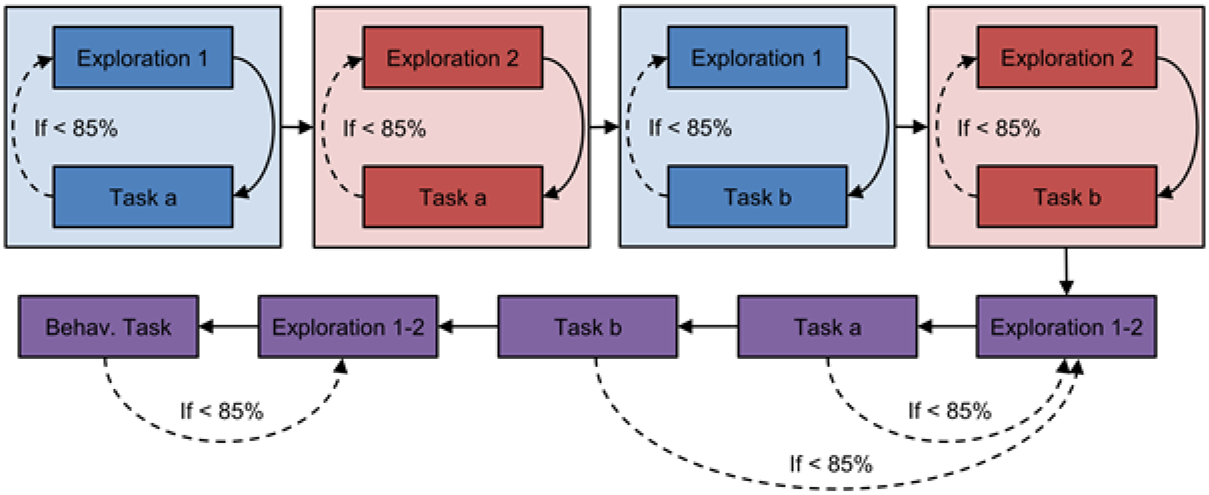


***Figure S2.*** *Flowchart depicting the structure of the online training session. Exploration 1 refers to the exploration of the two versions of one of the rooms, exploration 2 refers then to the exploration of the two versions of the other room, and Exploration 1-2 refers to the exploration of both versions of both rooms. For each task, if participants were not able to have an 85% accuracy, participants had to perform again the exploration.*

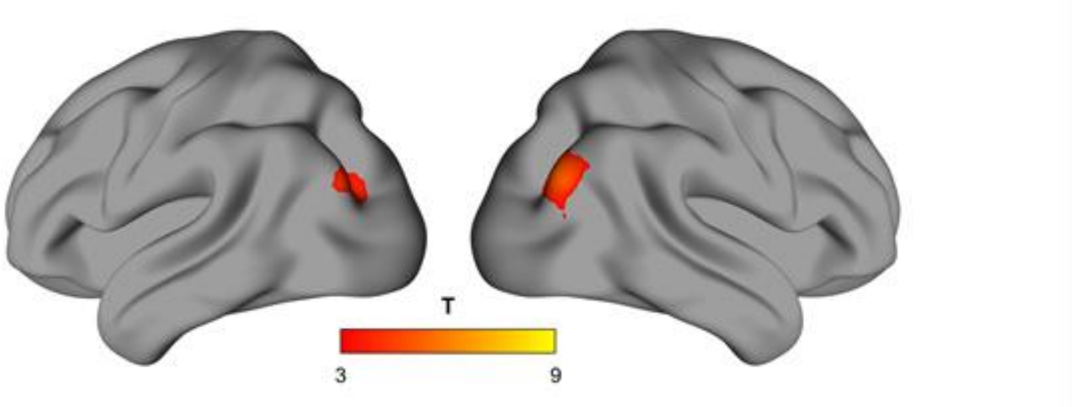


***Figure S3.*** *Whole-brain searchlight RSA during the reference window showed a bilateral activation of the occipital place area related to facing direction. A conventional cluster-extent-based inference threshold was used (voxel level at p* $< 0.001$*; cluster-extent FWE p* $< 0.05$*).*

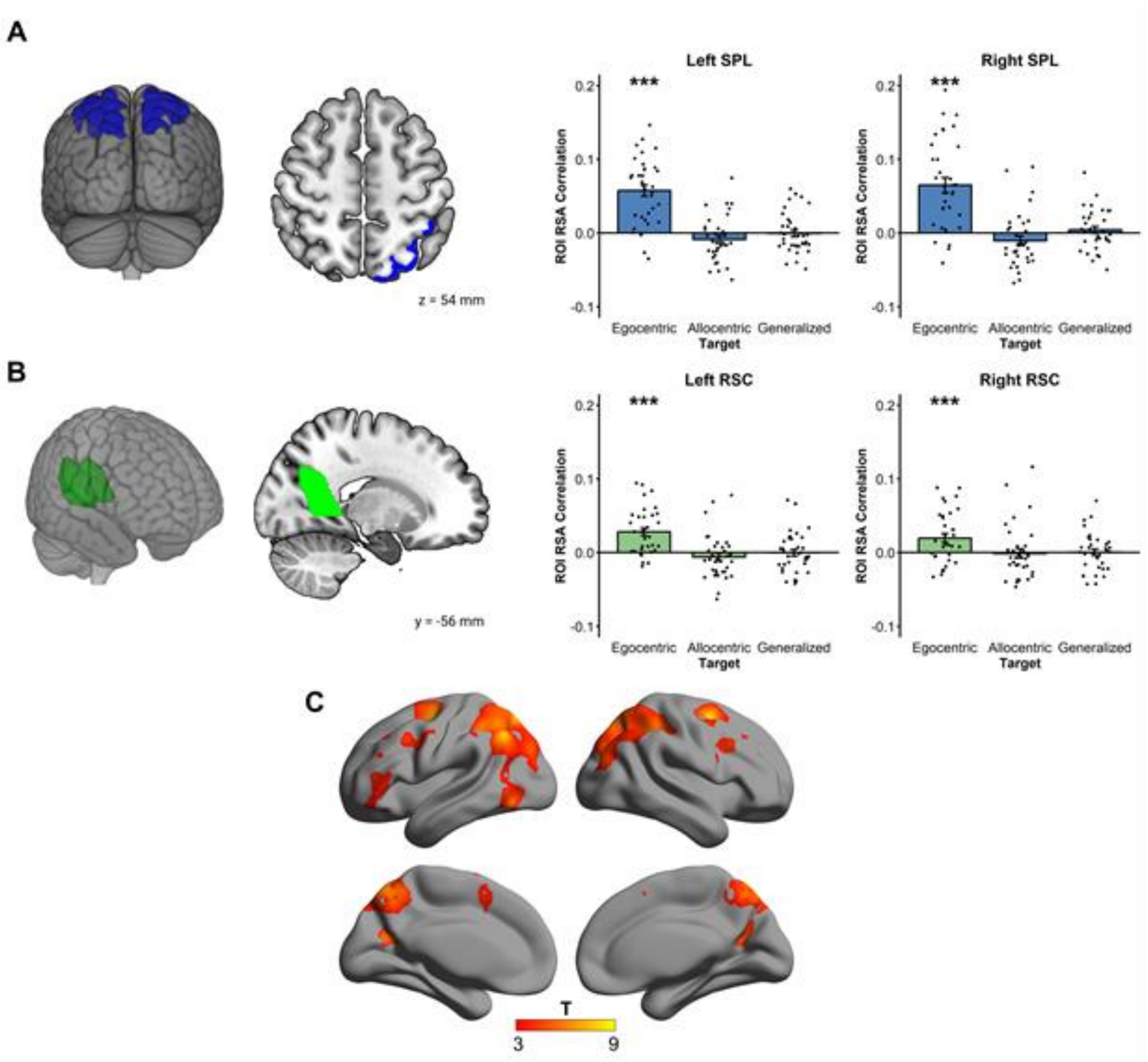


***Figure S4.*** *RSA Results for egocentric goal direction. (A) Both SPL showed reliable egocentric goal direction coding in the target window. (B) Both RSC showed reliable egocentric goal direction coding in the target window. (C) Whole-brain searchlight RSA confirmed the large parietal involvement in representing egocentric goal direction, showing further activations in the dorsal premotor area, the left posterior middle frontal gyrus, the left posterior cingulate cortex, and the left pars triangularis. (*** $p < .001$; voxel level at $p < 0.001$; cluster-extent FWE $p < 0.05$).*

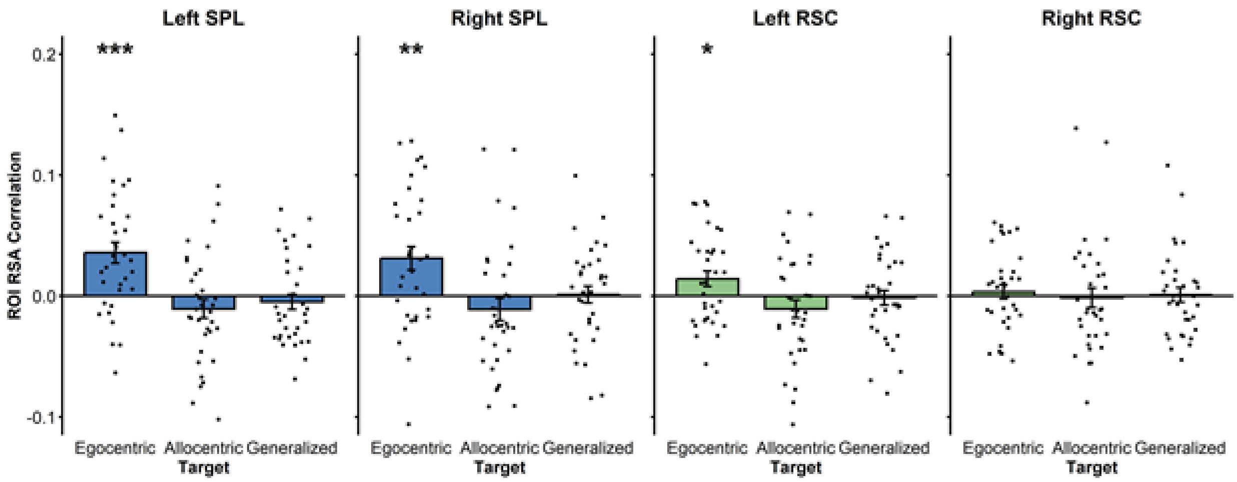


***Figure S5.*** *ROI analyses of the egocentric target direction in the parietal ROIs excluding the Front direction and controlling for response times (* $p < .05$*, **$p < .01$*, ***$p < .001$*).*

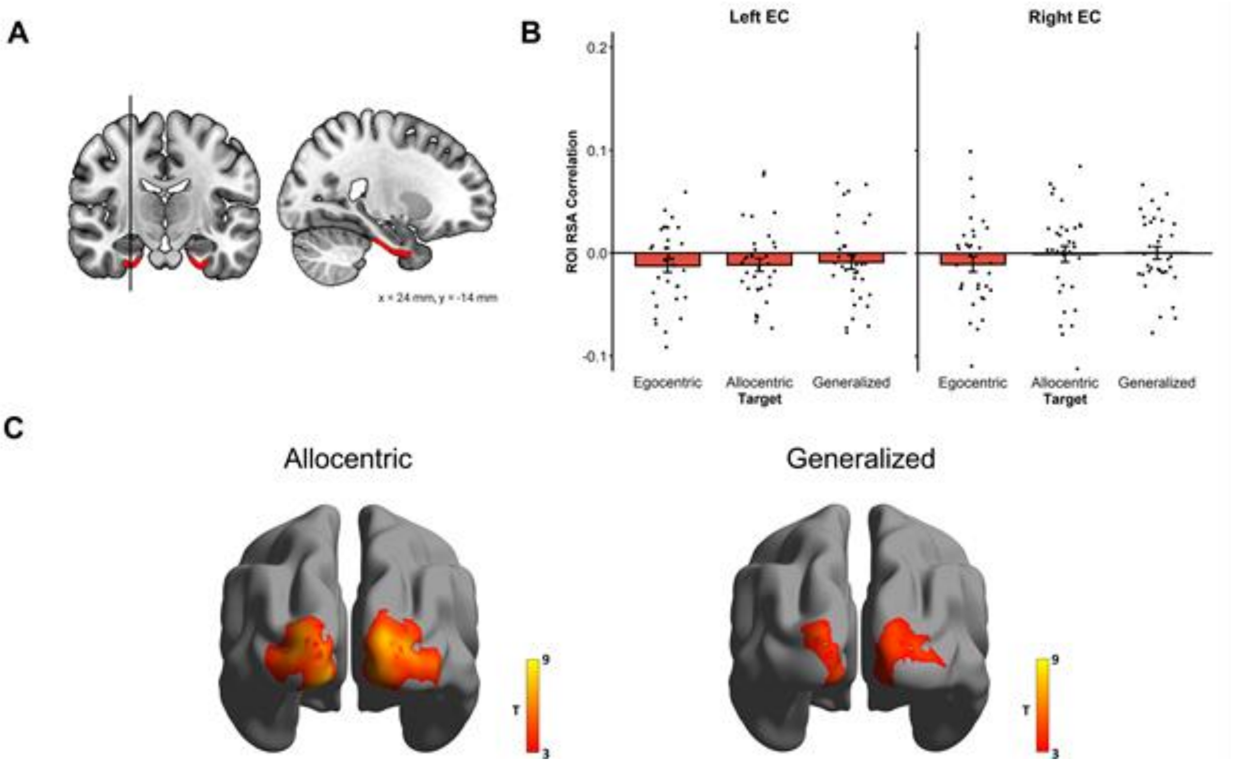


***Figure S6.*** *RSA Results for the allocentric goal direction. (A) EC ROIs (B) No reliable allocentric nor egocentric coding was observed in the ECs. (C) Allocentric and Allocentric-generalized whole-brain searchlight RSA results.*

| Region | MNI coordinates | | | Cluster size ($mm^3$) | t | $p_{FWE}$ |
|---|---|---|---|---|---|---|
| | x | y | z | | | |
| L angular gyrus | -48 | -62 | 44 | 103680 | 8.17 | < .001 |
| L dorsal premotor area | -26 | -8 | 58 | 11776 | 7.56 | .002 |
| L posterior middle frontal gyrus | -46 | 4 | 36 | 5112 | 4.96 | .008 |
| L posterior cingulate cortex | -16 | -62 | 20 | 3136 | 6.80 | .02 |
| L pars triangularis | -48 | 40 | 8 | 6344 | 4.51 | .006 |
| R dorsal premotor area | 24 | 0 | 54 | 8192 | 8.98 | .004 |
| R posterior middle frontal gyrus | 46 | 18 | 32 | 5544 | 5.54 | .002 |

***Table S1.*** *Results of the egocentric whole-brain analyses. A conventional cluster-extent-based inference threshold was used (voxel level at $p < 0.001$; cluster-extent FWE $p < 0.05$).*

| Region | MNI coordinates | | | Cluster size ($mm^3$) | t | $p_{FWE}$ |
|---|---|---|---|---|---|---|
| | x | y | z | | | |
| **Allocentric** | | | | | | |
| L occipital cortex | -16 | -88 | -12 | 17088 | 8.36 | < .001 |
| R occipital cortex | 18 | -98 | 10 | 19928 | 5.54 | < .001 |
| **Generalized** | | | | | | |
| L occipital cortex | -20 | -98 | 12 | 6128 | 5.23 | .003 |
| R occipital cortex | 16 | -96 | 12 | 9800 | 5.27 | .002 |

***Table S2.*** *Results of the allocentric and allocentric generalized goal whole-brain analyses. A conventional cluster-extent-based inference threshold was used (voxel level at $p < 0.001$; cluster-extent FWE $p < 0.05$).*
